## Supplemental document for "Sigma Oscillations Protect or Reinstate Motor Memory Depending on their Temporal Coordination with Slow Waves"

### Supplementary material & methods

#### Sample size estimation

The sample size was determined with a power analysis based on the paper of Cousins et al. **(1)** which reports, to our knowledge, the closest paradigm to the present one in the motor memory domain. This power analysis was performed through the G*Power software **(2)**. Specifically, we extracted the *Cohen's d* **(3)** effect size from the reported means in Cousins et al. ($d$ = 0.54) and converted into an $f$ effect size to perform the required ANOVA power analysis ($f$ = 0.26). Yet, to be more conservative, we set this effect size to 0.2. The average correlation coefficient was set at r = 0.8 between repeated measures and the sphericity correction was set to 1. The dependent variable was the offline changes in performance speed (see methods for details). These gains were computed for two time points: post-nap and post-night. The primary contrast of interest was a reactivation Condition (2 levels: reactivated sequence vs. non-reactivated sequence) by Time-point (post-nap vs. post-night) interaction tested with a repeated measure ANOVA (rmANOVA). The effect reaches a power of 95.9% with 24 participants at an alpha error probability of 0.05.

#### Vigilance

Vigilance was tested with a one-way rmANOVAs on both the median RTs of the PVT and the Stanford Sleepiness Scale score with Session as 3-level factor (pre-nap, post-nap, and post-night). The results presented in Table S2 indicate that vigilance did not differ among sessions.

#### Random SRTT

To highlight that improvement in movement speed was specific to the learned sequences as opposed to general improvement of motor execution, we computed the *overall performance gain* for both the sequential SRTT (first 4 blocks of t pre-nap raining vs. 4 last blocks of post-night training collapsed across reactivated and non-reactivated sequences) and the pseudo-random version of the SRTT (4 blocks pre-nap session vs. 4 blocks post-night session). We tested whether the overall performance gains on the learned sequences were larger than on the pseudo-random version of the SRTT using a two-tailed paired Student t-test (see supplemental results).

#### MSL/Sequential SRTT

As exploratory analysis, we ran a two-way rmANOVA with Condition (reactivated vs. non-reactivated) and Block (16 blocks of post-night training) as within-subject factors on the mean RT in each block of the Post-night training session of the sequential SSRT in order to assess whether any effects of TMR were sustained across the 16 blocks of the post-night retest.

#### Generation task

This task aimed at testing participants' explicit knowledge of the sequences as well as the strength of the association between the sequences and their corresponding auditory cues. During the generation task, participants were presented with the auditory cues specific to the learned sequences and were instructed to self-generate the corresponding motor sequences. Participants completed 4 consecutive attempts for each cue / sequence pair. The order of the pairs was randomized. Accuracy was emphasized during this task. A trial was classified as “correct” if the key pressed by the participant was in the correct ordinal position with respect to the sequence acoustically cued. The percentage of correct ordinal positions was computed per sequence and per attempt. The generation accuracy per sequence was computed by averaging across attempts for each time point separately (pre-nap and post-night sessions). We tested whether generation accuracy of the reactivated sequence during the pre-nap generation task was correlated (Pearson’s correlation) to the TMR index (see supplemental results).

### Supplementary results

#### Cue distribution

As an exploratory negative control analysis, the number of the different auditory cues (associated vs. unassociated) in the different sleep stages (Wake vs NREM1 vs NREM2 vs NREM3 vs REM) was compared with an rmAnova using the number of auditory cues as dependent variable, cue type as 2-level factor, sleep stage as 5-level factor, and participant as repeated measure. The number of the cues was different between sleep stages as expected since stimulation was restricted to NREM2-3 stages (main effect of Stage: $F(4,92) = 22.18$; p-value = 7.68e-06; $\eta^2 = 0.49$). Yet, the number of cues did not differ between Cue Type (main effect of Cue: $F(1,23) = 1.67$; p-value = 0.21 or a Stage by Cue interaction: $F(4,92) = 0.21$; p-value = 0.93).

#### Random SRTT

To determine whether the increase in performance speed on the SRTT was due to *motor sequence learning* rather than a mere increase of motor execution, we tested whether performance improvement on the learned sequences was larger than on the pseudo-random version of the SRTT (Figure 2 in main manuscript). We thus compared (two-tailed paired Student t-test) the overall gains in performance between the two SRTT types. Overall performance gains were significantly higher for the sequential SRTT as compared to the random SRTT ($t = 21.69$, $df = 23$, p-value < 2.2e-16; Cohen's $d = 4.43$). Thus, the RT decrease observed in the sequential SRTT reflect motor sequence learning rather than a mere improvement in motor execution.

#### Baseline performance

The baseline performance speed (i.e. pre-nap) was compared between sequences type (sequence A vs. sequence B) regardless the reactivation condition. Two two-way rmANOVAs with Sequence (sequence A vs. sequence B) and Block ($1^{st}$ rmANOVA on the 16 blocks of the pre-nap **training** and $2^{nd}$ rmANOVA on the 4 blocks of the pre-nap **test**) as within-subject factors was performed on performance speed. The performance during the pre-nap training was not significantly different between the sequence A and the sequence B (main of effect Sequence: $F(1,23) = 2.18$; p-value = 0.15). The RT significantly decreased across blocks (Main effect of Block: $F(15,345) = 34.82$; p-value = 2.04e-26; $\eta^2 = 0.6$). However, this decrease was not different between the two sequences (Block by Sequence interaction: $F(15, 345)= 1.11$; p-value = 0.35). During the pre-nap test, the main effect of Block was significant ($\eta^2 = 0.17$ and $F(3,69) = 6.67$; p-value = 0.001; $\eta^2 = 0.22$) as well as the interaction between the Block and the Sequence ($F(3,69)= 3.3$; p-value = 0.025; $\eta^2 = 0.13$). The main effect of sequence was not significant although very close from the alpha threshold ($F(3,69)= 4.11$; p-value = 0.054). After removal of the first block of the pre-nap test showing non-plateau performance (see main text), the main effect of Sequence was not significant ($F(1,23) = 0.99$; p-value = 0.33) nor was the block effect ($F(2,46) = 1.56$; p-value = 0.22). Yet, the interaction between the Block and the Sequence remained significant ($F(2,46)= 3.37$; p-value = 0.043; $\eta^2 = 0.13$). However, as reactivation conditions were counterbalanced between sequences A and B, this small effect is not considered as confounding to test our hypothesis.

In terms of accuracy, baseline performance (i.e. pre-nap) was compared between the two reactivation conditions. Two two-way rmANOVAs with Condition (reactivated vs. non-reactivated) and Block ($1^{st}$ rmANOVA on the 16 blocks of the pre-nap training and $2^{nd}$ rmANOVA on the 4 blocks of the pre-nap test) as within-subject factors was performed on the averaged accuracy in each block of sequential SRTT. In the pre-nap training, there were no main effects (Condition: $F(1,23) = 0.74$; p-value = 0.4; Block: $F(15,345) = 1.3$; p-value = 0.2) nor interaction ($F(15,345) = 0.39$; p-value = 0.98). This was also the case for the pre-nap test (Condition: $F(1,23) = 1.01$; p-value = 0.75; Block: $F(3,69) = 2.24$; p-value = 0.88; Interaction: $F(3,69) = 1.06$; p-value = 0.37). When comparing the baseline accuracy performance between sequence type (i.e., sequence A vs. Sequence B), the rmANOVA performed on the pre-nap training highlighted a main effect of Sequence ($F(1,23) = 7.28$; p-value = 0.01; $\eta^2 = 0.24$), but no significant effect of the Block factor nor interaction ($F(15,345) = 1.3$; p-value = 0.2 and $F(15,345) = 1.03$; p-value = 0.43 respectively). The sequence effect was not significant during the pre-nap test and no block effect or block by sequence interaction were observed (Sequence: $F(1,23) = 0.25$; p-value = 0.62; Block: $F(3,69) = 0.22$; p-value = 0.88; Interaction: $F(3,69) = 0.41$; p-value = 0.75).

##### Offline gains in performance accuracy

A rmANOVA with Time-point (post-nap vs. post-night) and Condition (reactivated vs. non-reactivated) was performed on the offline gains in performance accuracy. The offline gains were significantly higher at the post-night time-point as compared to the post-nap time point ($F(1,23) = 18.84$; p-value = 0.0002; $\eta^2 = 0.91$). Yet, the offline gains in performance accuracy were not different between the reactivated and the non-reactivated sequences ($F(1,23) = 0.54$; p-value = 0.47). The interaction between the Condition and the Time-point was also not significant ($F(1,23) = 0.2$; p-value = 0.66).

##### RTs along the post-night training

In order to assess whether any effects of TMR were sustained across the post-night retest session (see Figure 2, main manuscript), an exploratory two-way rmANOVA was performed on performance speed during the Post-night training session with Condition (reactivated vs. non-reactivated) and Block (16) as within-subject factors. There was no interaction between the Condition and Block ($F(15, 345) = 1.05$; p-value = 0.4). The RT significantly decreased with blocks ($F(15,345) = 9.94$; p-value = 6.96e-17; $\eta^2 = 0.3$), highlighting that performance still improved during this next-day training session. Finally, the main effect of Condition was marginally significant ($F(1,23) = 3.37$; p-value = 0.08; $\eta^2 = 0.13$), hinting towards a sustained effect of TMR on memory consolidation.

##### Generation task

The results on the generation task (Table S3) were not significantly correlated with the TMR index ($r = 0.25$, $t = 1.22$, df = 22, p-value = 0.24).

##### ERP

In order to identify specific time windows to compare ERP amplitude between conditions at the peak and the trough of the potential, we used CBP approaches on ERP data computed across conditions (see methods in the main text). Results show that across conditions, ERP amplitude was significantly different from zero between 0.44 – 0.63 sec at the trough (alpha threshold = 0.025, cluster p-value = 0.044; *Cohen's d* = -0.56) of the potential (see supplementary Figure S2). The effect reported in the main manuscript across channels is more pronounced on the Frontal (Fz) and the Central (Cz, C3, and C4) electrodes (see Figure S3).

#### Correlation analyses between TMR and EEG-derived metrics

The correlations were computed using One-sided Spearman[#] tests (supplemental Table S1.6) and were corrected for 3 multiple comparisons. The TMR index was not significantly correlated with: (1) the relative change between the density of spontaneous SO during the associated and unassociated cue stimulation (S = 2486, p-value = 0.97); (2) the relative change between the density of spontaneous spindles during the associated and unassociated cue stimulation (S = 1412, p-value = 0.24); nor (3) the relative change between the amplitude of the negative peak of the ERP following the associated and unassociated auditory cues (S = 2282, p-value = 0.73).

### Supplementary tables

**Table S1**. List of the deviations from the pre-registered analyses followed by their justification. These deviations are marked with a # in the main manuscript.

| | Pre-registered | Final report |
|---|---|---|
| 1 | Only right-handed participants | Both right- and **left**-handed participants |
| | Justification: Based on the bimanual nature of our motor task, we elected to not restrict our participant pool to only right-handers in order to facilitate recruitment (N= 2 left-handers). | |
| 2 | Pre-nap performance for offline gain computation on last 4 blocks of the MSL | Pre-nap performance for offline gain computation on last **3** blocks of the MSL |
| | Justification: See main manuscript for details. Briefly, against our expectations based on previous research using learning of a single sequence, participants only reached plateau performance on the two sequences starting on block 2 of the pre-nap test. In order to meet the performance plateau pre-requisite to compute offline gains in performance, we therefore excluded the first block of the pre-nap test and computed offline gains based on the last 3 blocks of the pre-nap test which showed stable performance levels for both sequences. | |
| 3 | We will classify auditory evoked responses into evoked SO if the standard criteria of a SO are met (negative peak ≤ -40 µV and the peak-to-peak amplitude ≥ 75 µV). Mean auditory-evoked SO amplitude will be computed for each subject in each condition separately. | Auditory-evoked responses were averaged **across all trials** for each condition. Mean auditory **ERP** amplitude was computed for each subject in each condition separately. |
| | Justification: The number of auditory ERPs reaching the pre-registered amplitude criteria was not sufficient to perform a powerful statistical analysis. This issue being highlighted in previous research (4), we followed similar procedures and averaged all the auditory-evoked responses (irrespective of their amplitude) on the one hand and all the detected SO on the other hand (4). | |
| 4 | To analyze spindle activity, we will compute Time-Frequency representations (TFR) using Morlet Wavelet decomposition with a width of five cycles per wavelet (m=7) at center frequencies between 12 and 16 Hz (sigma frequency band), in steps of 0.5 Hz and 10 ms. | To analyze spindle activity, we computed Time-Frequency representations (TFR) using **an adaptive sliding time window of five cycles length per frequency ($\Delta t = 5/f$; 20-ms step size), and estimated power using the Hanning taper/FFT approach between 5 and 30 Hz**. |
| | Justification: For completeness, we decided to broaden our analysis from 12-16 Hz to 5-30 Hz. Consequently, the Morlet wavelet decomposition approach was not appropriate to capture low (around 5 Hz) frequency band in regards of the epoch size. Note that any effects observed outside the pre-registered frequency band (i.e, sigma) are reported in the main text as the result of exploratory analyses. A similar frequency window approach was used for the PAC analyses. | |
| 5 | The mean of the circular angle values for SO-spindle coupling will be compared between stimulation blocks of associated vs. unassociated auditory cues using a one-tailed paired student t test. | The mean of the circular angle values for SO-spindle coupling was compared between stimulation blocks of associated vs. unassociated auditory cues using **a Watson's goodness of fit test for the von Mises or circular uniform distribution.** |
| | Justification: This statistical test was more appropriate for circular data as compared to the classical student t test (5). | |
| 6 | Parametric statistical testing will be applied, namely Student t-tests for comparison of means and Pearson tests for correlation analyses. | We performed non-parametric statistical testing whenever the normality criterion was not met, **namely Wilcoxon signed-rank tests for comparison of means and Spearman tests for correlation analyses.** |
| | Justification: The assumption of normality was not met for some metrics; we thus used the equivalent non-parametric test in these cases. | |
| 7 | Pearson correlations will be performed between the TMR index and the relative change between | The correlation between TMR index and the difference between sigma band power time-locked |

| | | |
|---|---|---|
| | sigma band power time-locked to the associated auditory cues and the sigma band power time-locked to unassociated auditory cues. | to the associated auditory cues and the sigma band power time-locked to unassociated auditory cues was performed using **CBP tests due to the time-frequency dimension of the data.** |
| | Justification: Correlation analyses between the TMR index and the raw difference in sigma band power time-locked to the associated and unassociated auditory cues were performed using Cluster Based permutation tests (6). This approach was preferred as it is more conservative than the analysis initially pre-registered and allows to handle and correct for the large time-frequency dimension of the data. Accordingly, CBP approaches were also used for the correlation analyses between PAC and TMR index. | |

**Table S2**. Group averaged (+/- 95% confidence interval) sleep characteristics leading up to the experimental session and vigilance assessments at time of testing

| | |
|---|---|
| **Sleep duration[a]** | |
| Mean across the 3 nights (minutes) | |
| Night 1 | 481.5 [461.5 – 501.5] |
| Night 2 | 488.2 [471.2 – 505.2] |
| Night 3 | 502 [482.1 – 521.8] |
| **St. Mary's questionnaire on Night 3 quality** | |
| Quality | 4.7 [4.3 – 5.1] |
| Duration (minutes) | 443.5 [423.3 – 463.8] |
| **Psychomotor Vigilance Task[b]** | |
| Pre-nap | 300.2 ms [289.7 –310.6] |
| Post-nap | 297.5 ms [288.9 – 306] |
| Post-night | 294.7 ms [285.1 – 304.2] |
| One-way rmANOVA results | $F(2,46)=1.47$; p-value = 0.24 |
| **Stanford sleepiness score** | |
| Pre-nap Session | 2.4 [2.1 – 2.7] |
| Post-nap Session | 2.7 [2.3 – 3.1] |
| Post-night Session | 2.3 [1.9 – 2.7] |
| One-way rmANOVA results | $F(2,46)=1.69$; p-value = 0.2 |

*Notes*. Values are means [lower and upper limit of the 95% Confidence Interval - CI]. REM: Rapid Eye Movement. [a] Sleep duration was computed as the mean across the actigraphy data and the sleep diary for the three nights before the experimental day. [b] Median of reaction times computed across the 100 trials of each session.

**Table S3**. Group averaged [lower and upper limit of the 95% CI] accuracy for the generation task

| Pre-nap | | Post-night | |
|---|---|---|---|
| Reactivated | Non-reactivated | Reactivated | Non-reactivated |
| 78.77 [67.51 – 90.03] | 69.4 [57.94 – 80.87] | 74.74 [61.88 – 87.6] | 81 [69.73 – 92.26] |

**Table S4**. Mean number [lower and upper limit of the 95% CI] of sleep events across participants detected at the single channel level per condition of blocks (either stimulation associated/unassociated or rest).

**Slow waves**

| | Associated | Unassociated | Rest | Entire Nap |
|---|---|---|---|---|
| **FZ** | 159.7 [80.3 – 239.1] | 161 [83.3 – 238.8] | 155.5 [88.9 – 222.1] | 469.5 [255.6 - 683.4] |
| **CZ** | 139 [71.9 – 206.2] | 139.4 [74.5 204.3] | 139 [80.8 – 197.2] | 411.6 [231.4 - 591.7] |
| **PZ** | 121.3 [58.8 – 183.9] | 115.7 [59.2 – 172.2] | 109.7 [60.9 – 158.7] | 336.9 [178.7 - 495.1] |
| **OZ** | 52.6 [18.5 – 86.7] | 51.6 [19.8 – 83.5] | 42.8 [20.9 – 64.6] | 130.5 [52.7 - 208.3] |
| **C3** | 99.8 [41.3 - 158.3] | 97.2 [41.9 – 152.6] | 91.3 [47.3 – 135.4] | 284.3 [132 - 436.7] |
| **C4** | 107.6 [46.2 – 169.1] | 102.2 [46.1 158.2] | 100.5 [52.7 – 148.4] | 297.4 [141.4 - 453.4] |

**Spindles**

| | Associated | Unassociated | Rest | Entire Nap |
|---|---|---|---|---|
| **FZ** | 27.7 [14.6 – 40.8] | 31.9 [17.3 – 46.4] | 49.2 [30.7 – 67.6] | 102.6 [62.8 - 142.4] |
| **CZ** | 28.4 [14.9 – 41.9] | 34.3 [19.8 – 48.9] | 54 [36 – 71.9] | 114.1 [72.5 - 155.7] |
| **PZ** | 27.5 [14.8 – 40.1] | 34.2 [20.2 – 48.3] | 59 [42 – 75.9] | 119.5 [80.6 - 158.5] |
| **OZ** | 11.8 [5 – 18.6] | 11.2 [6 – 16.4] | 22.7 [12.1 – 33.3] | 37.5 [19.7 - 55.4] |
| **C3** | 25 [13.1 - 37] | 30.6 [16 – 45.2] | 52.4 [34.9 – 69.9] | 105.7 [66.5 - 144.9] |
| **C4** | 27 [12.6 – 41.5] | 31 [16.3 – 45.7] | 54.1 [34.8 – 73.3] | 108.5 [65 - 152.1] |

#### Supplementary figures

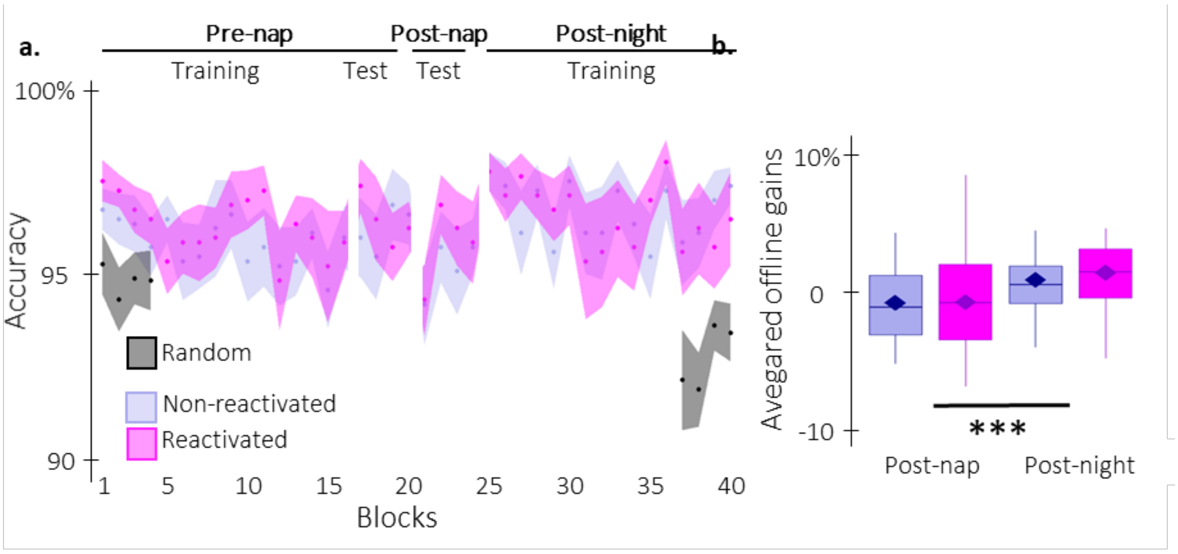


*Figure S1: Behavioral results on performance accuracy. a. Performance accuracy* (*mean percentage of* correct responses) *across participants plotted as a function of blocks of practice during the pre and post-nap sessions (+/- standard error) for the reactivated (magenta) and the non-reactivated (blue) sequences.* ***b. TMR effect****. Offline gains in performance accuracy averaged across participants (box: median and first(third) as lower(upper) limits; whiskers: 1.5 x InterQuartile Range (IQR)) for post-nap and post-night time-points and for reactivated (magenta) and non-reactivated (blue) sequences. ***: p-value < 0.001*

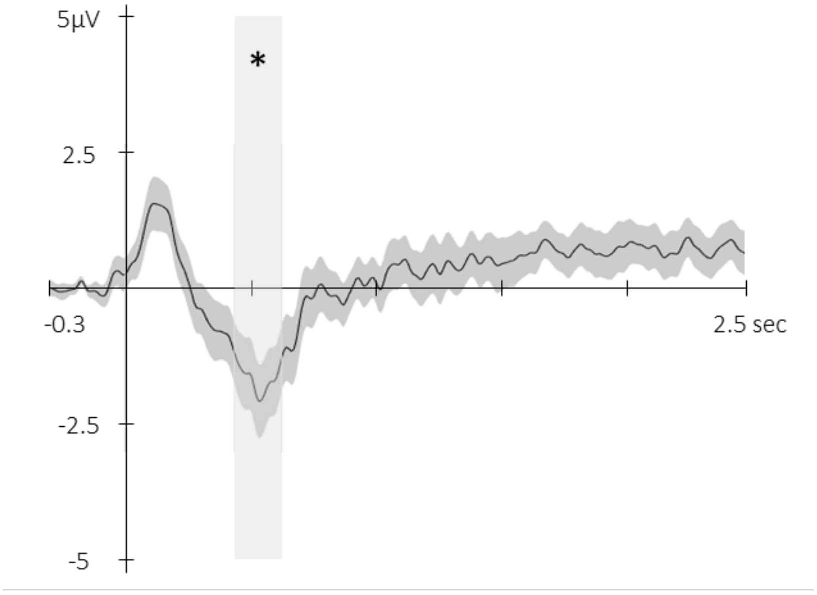


*Figure S2: Event-related Potential across conditions.* *Potentials averaged across all participants and all EEG channels (+/- standard error) evoked by the auditory cue regardless of its type from -0.3 to 2.5 sec relative cue onset. The grey region represents the temporal window (trough) in which ERPs across conditions were significantly different from zero. *: p-value < 0.05*

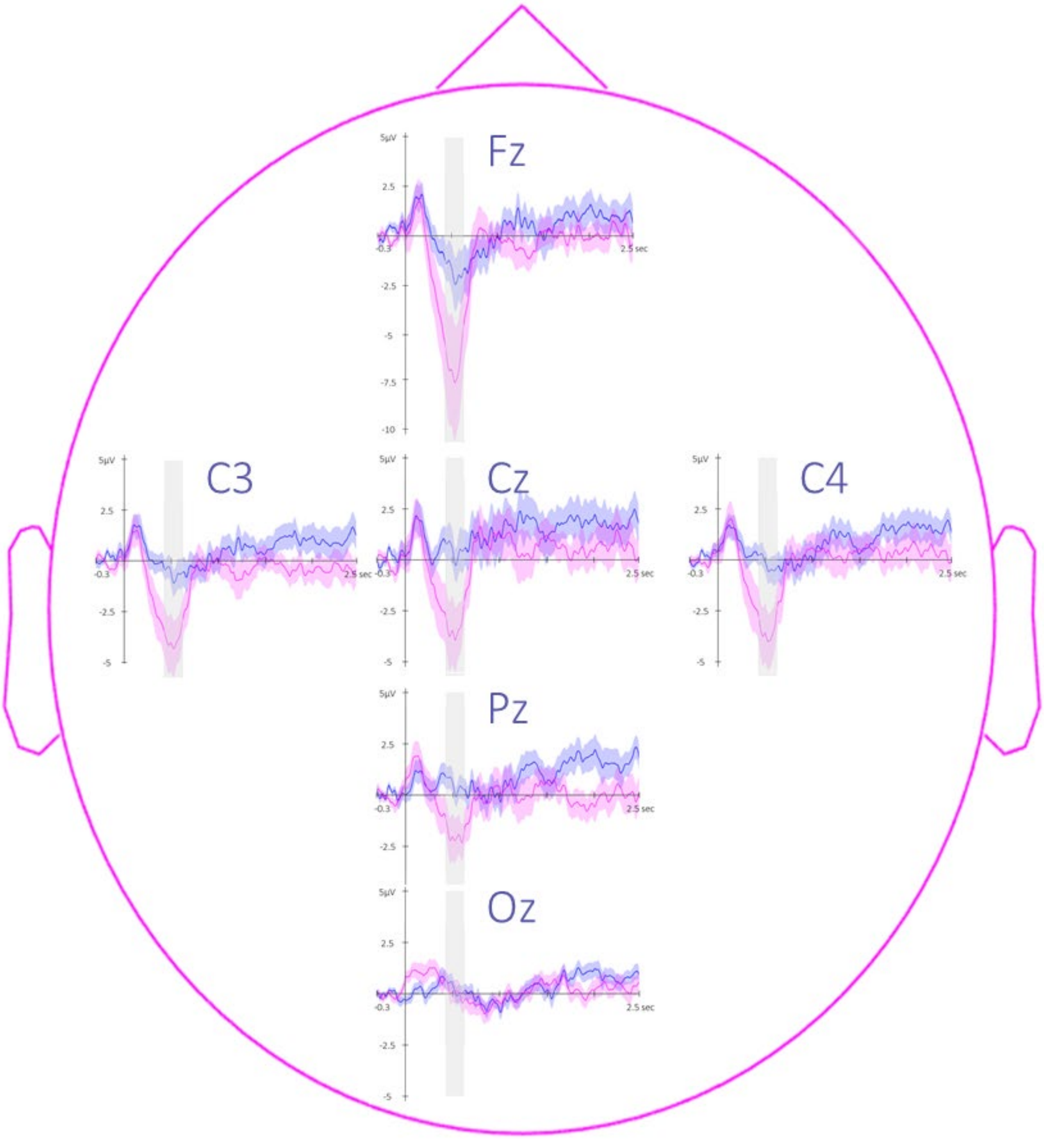


*Figure S3: Topography of Event-related Potentials.* *Potentials evoked by the associated (magenta) and the unassociated (blue) auditory cues from -0.3 to 2.5 sec relative to cue onset averaged across all participants (+/- standard error). Note that the effect reported in the main manuscript across channels is more pronounced on the Frontal (Fz) and the Central (Cz, C3, and C4) electrodes. The grey region represents the temporal window (trough) in which ERPs across conditions were significantly different from zero.*
